# Haematobiochemical Alterations and Lesion Characterization Induced by Haemonchosis in Sheep Slaughtered at Gondar ELFORA Abattoir, North West Ethiopia

**DOI:** 10.1101/2024.10.03.616054

**Authors:** Birhan Anagaw Malede, Mersha Chanie Kebede, Asnakew Mulaw Berihun, Muluken Yayeh Mekonnen, Mohammed Yesuf, Tadegegn Mitiku, Mastewal Birhan, Ambaye Kenubih, Abraham Belete Temesgen

## Abstract

Haemonchosis is a major gastrointestinal parasitic infection in sheep caused by *Haemonchus contortus*. An abattoir-based cross-sectional study was conducted from January to September 2024 to assess the Haematobiochemical alterations and lesion characterization induced by Haemonchosis in sheep slaughtered at the Gondar ELFORA abattoir. The study involved 60 male local breed sheep, divided into 30 infected and 30 non-infected controls. The selection process involved postmortem examination of the abomasum tissues, incision, palpation, and visual inspection. Blood samples were taken for hematology and serum biochemical profiles, and a three cm tissue sample with typical *Haemonchus contortus* lesions was also taken. The study found significant reductions in hemoglobin, hematocrit, corpuscular volume, hemoglobin, and red blood cell counts in the infected group compared to the non-infected group. However, white blood cells, monocytes, neutrophils, and eosinophils were significantly higher in the infected group. Biochemical parameters showed significant reductions in total protein, albumin, globulin, and albumin to globulin ratio in the infected group. Erythrocyte indices indicated microcytic normochromic anemia. Gross examination revealed hemorrhages, dark brown abomasal contents, blood streaks, and a nodular lesion. Microscopic analysis revealed tissue-dwelling worms, submucosal hemorrhage, mucosal gland hyperplasia, thickened muscularis, and hyperplastic abomasal glands. The alterations in haematobiochemical parameters support the findings from gross and microscopic lesions. Thus, integrating haematobiochemical analysis with gross and microscopic lesion characterization improves the diagnosis of haemonchosis. Due to hypoproteinemia observed, it is advisable to supplement helminth-infected animals with protein-rich feeds, such as legumes.

## 1. INTRODUCTION

Sheep are an important part of the livestock industry in developing nations like Ethiopia. They provide a substantial amount of income for farmers, dealers, and the state because they are mostly raised in rural areas, sold locally as organic meat, and used in religious festivals [1, 2, 43]. Increased purchasing power, expanding international markets, and population growth are the main drivers of the growing demand for sheep. Sheep are prized for their meat, wool, and skin; the foreign exchange profits from these products support local and global economies. Dairy products and sheep milk also present market opportunities and possible health benefits [45]. However, there are significant health issues facing the industry, mainly related to gastrointestinal parasites like the *Haemonchus species*, which reduce productivity, raise mortality, and require expensive treatment [13].

*Haemonchus contortus* is the parasitic cause of hemochosis, a serious gastrointestinal disease that affects sheep [33]. The highly pathogenic blood-sucking parasite *Haemonchus contortus*, commonly referred to as the “barber’s pole worm,” lives in the abomasum of small ruminants and has the potential to cause infections that could be fatal [51]. Nematodes such as *Haemonchus, Ostertagia, Teladorsagia*, and *Trichostrongylus* rely on the abomasum as a vital location [17, 58]. Hemonchosis can present in three different ways: hyperacute, acute, and chronic. Each type of hemochosis has a different clinical manifestation and range of severity [11, 37].

It has been demonstrated that *Haemonchus contortus* infection results in important hematological and biochemical alterations. Erythrocyte counts, hemoglobin concentrations, and hematocrit (Hct) all significantly decline as a result, suggesting anemia. White blood cell (WBC) counts and eosinophil levels also rise significantly as a result of the infection, indicating an immune reaction to the parasite. According to biochemical analysis, *H. contortus* infection also results in a marked reduction in calcium, iron, and total protein levels, which may worsen the animals’ health [4, 26, 47]. A great number of viable, actively moving adult *Haemonchus contortus* parasites, pale mucous membranes, fluid-filled abomasal contents stained dark brown with bloody flakes, and ecchymotic hemorrhagic and hyperemic mucosal folds with focal ulcerations would all be grossly visible. Histopathologically, the mucosa and stomach glands would show hemorrhages, edema, lymphocytic and extensive eosinophilic infiltration, blood vessel congestion, and desquamation of the abomasal villi [23, 35].

One important link in the livestock supply chain, the Gondar ELFORA abattoir, deals with a significant amount of small ruminants. For the purpose of putting targeted control measures into place and guaranteeing the production of safe and healthful meat products, it is essential to comprehend the effects of *Haemonchus contortus* infection on animals processed at this abattoir [45]. An invaluable resource for learning about disease epidemiology is the butcher shop. Both antemortem and postmortem examinations are performed during meat inspection at slaughterhouses; post-mortem inspection is crucial for assessing clinical indicators and pathological processes that impact the quality of the meat [29].

Despite wide research by international scholars documenting the hematobiochemical and pathological effects of *Haemonchus contortus* in sheep [3, 23, 54, 58], there remains limited research conducted within Ethiopia [1]. Therefore, the present study was conducted to compare the hematological and biochemical profiles of *Haemonchus contortus*-infected and non-infected sheep and to characterize the pathological lesions associated with this parasite infection.

## 2. MATERIALS AND METHODS

### 2.1. Study Area and Animal

The study was carried out at the privately owned Gondar ELFORA Abattoir, which is located in Gondar, the capital of the Central Gondar Administrative Zone in the Amhara Regional State of Ethiopia, from January to September of 2024. Gondar is situated at geographic coordinates 12°36′ N latitude and 37°28′ E longitude. It is roughly 727 kilometers from Addis Ababa and sits at an elevation of 1,800 to 2,220 meters (Figure 1). Weekly, the abattoir processes twenty to twenty-five sheep for the nearby market. March through May is the shortest rainy season in the region; June through September is the longest. There are 1,936,514 cattle, 524,083 sheep, 682,264 goats, 36,828 horses, 12,473 mules, 223,116 donkeys, and 3,165,068 chickens in the area [56]. The sheep that were killed at this location came from different Central Gondar zone districts. Male sheep of the local breed that were brought to the slaughterhouse for human consumption were the subject of the study. Raised in a vast farming system, these sheep were driven to the abattoir by truck and kept in lairage for a day prior to processing.

### 2.2. Study Design and Period

An abattoir-based cross-sectional study was conducted from January to September 2024 to assess the Haematobiochemical alterations and lesion characterization induced by Haemonchosis in sheep slaughtered at the Gondar ELFORA abattoir.

### 2.3. Sample Size and Sampling Method

A total of sixty male local breed sheep were purposively selected for this study, divided equally into two groups: 30 sheep infected with *Haemonchus contortus* and 30 non-infected controls. The selection process was based on the confirmed presence of *H. contortus* in the abomasum, as determined through postmortem examination that involved systematic incision, palpation, and visual inspection of the abomasal tissues.

### 2.4. Criteria for Inclusion and Exclusion of Parasites

The study included abomasal tissues containing *Haemonchus contortus* parasites, while tissues with other parasites or mixed infections were excluded.

### 2.5. Sample Collection

A total of 60 blood samples were taken using 5 ml EDTA-coated vacutainers for hematological analysis and 5 ml plain tubes for serum biochemical evaluations. Additionally, a total of 20 representative abomasal tissue samples were obtained for histopathological examination from 30 infected sheep. Every sample had typical haemonchosis lesions and was roughly 3 cm in size. Finally, each sample was carefully sent to the University of Gondar’s Veterinary Pathology Laboratory.

### 2.6. Sample Processing

#### 2.6.1. Haematology analysis

Hematological analysis was conducted using a Sysmex automated hematology analyzer according to the manufacturer’s guidelines. Various parameters, such as mean corpuscular volume (MCV), mean corpuscular hemoglobin (MCH), mean corpuscular hemoglobin concentration (MCHC), hematocrit, hemoglobin (HGB), total erythrocyte count (TEC), total leukocyte count (TLC), and differential leukocyte count (DLC), were measured.

#### 2.6.2. Biochemical analysis

Anticoagulant-free blood samples were left to clot in a slanted position for three to four hours in preparation for the biochemical analysis. After that, the serum was separated, kept in labeled Eppendorf tubes at −20°C, and examined in accordance with Yesuf et al.’s [60] instructions. Using standard commercial test kits from Human GmbH (Wiesbaden, Germany) and an automated clinical chemistry analyzer, serum concentrations of Total Protein, Albumin, Globulin, and the Albumin/Globulin ratio were measured in accordance with manufacturer’s instructions.

#### 2.6.3. Gross examination

The external and internal surfaces of the abomasum were thoroughly examined for any abnormalities during the gross examination, and the results were meticulously recorded. A three cm sample of tissue was taken, including sections of the infected Haemonchus tissue as well as healthy tissue. After that, the sample was preserved in 10% neutral buffered formalin for histopathological examination.

#### 2.6.3. Histopathological examination

Twenty abomasal tissues that were infected with *Haemonchus contortus* were fixed in 10% neutral buffered formalin, cleared with xylene, dehydrated in increasing grades of ethanol, impregnated, and embedded in paraffin. Hematoxylin and eosin (H&E) staining, tissue sectioning at 5 μm, and DPX mounting were performed on the tissues. A light microscope was used to examine the slides, with magnifications ranging from 10x to 100x [6].

### 2.7. Data Analysis

After being collected, the data were coded, checked, and arranged before being added to an Excel spreadsheet (Microsoft Office Excel 2010). STATA version 18 was used for the statistical analysis. A t-test was performed to evaluate significant differences between the haematological and biochemical parameters of samples infected with *Haemonchus contortus* and samples that were not. If there was a statistically significant difference, the p-value had to be less than 0.05. The means ± standard errors (SE) were used to express the results. Pictures depicting both microscopic and gross lesions were utilized to illustrate descriptive statistics.

## 3. RESULTS

### 3.1. Haematological Findings

The hematological analysis indicated significant mean reductions (P < 0.05) in hemoglobin, hematocrit, mean corpuscular volume, mean corpuscular hemoglobin, and red blood cell counts in the infected group (6.58 ± 0.13, 22.52 ± 0.32, 25.74 ± 0.54, 6.75 ± 0.10, and 6.029 ± 0.179, respectively) compared to the non-infected group (12.96 ± 0.22, 35.82 ± 0.68, 36.76 ± 1.85, 10.85 ± 0.36, and 12.58 ± 0.229, respectively). In contrast, the mean values of white blood cells, monocytes, neutrophils, and eosinophils were significantly higher in the infected group (14.60 ± 0.26, 7.51 ± 0.20, 51.57 ± 1.54, and 15.06 ± 0.35, respectively) than in the non-infected group (8.21 ± 0.35, 3.55 ± 0.16, 33.54 ± 0.91, and 6.72 ± 0.39, respectively). The mean corpuscular hemoglobin concentration and lymphocyte count were within normal ranges, with no significant differences between the groups. The mean of basophils in both groups showed no change. The hematological parameters for non-infected sheep were within the established reference ranges (P < 0.05) (Table 1).

**Table 1.** Hematological Parameters of *H. contortus*-Infected and Non-Infected Sheep.

| Parameters | Mean ± SE | | Normal Range | t-value | p-value |
| --- | --- | --- | --- | --- | --- |
|  | Infected (n=30) | Non-infected (n=30) |  |  |  |
| HGB (g/dL) * | 6.58± 0.13 | 12.96± 0.22 | 9- 15 | 24.92 | 0.000 |
| HCT (%) * | 22.52± 0.32 | 35.82± 0.68 | 25-45 | 15.55 | 0.000 |
| MCV (fL) * | 25.74±0.54 | 36.76±1.85 | 28-40 | 5.78 | 0.000 |
| MCH (pg) * | 6.75± 0.10 | 10.85± 0.36 | 8-12 | 10.94 | 0.000 |
| MCHC (%) <sup>NS</sup> | 33.95± 6.82 | 32.14± 0.27 | 31-35 | 0.26 | 0.793 |
| RBC ($\times 10^6/\mu\text{L}$) * | 6.029±0.179 | 12.58± 0.229 | 9-15 | 23.23 | 0.000 |
| WBC ($\times 10^3/\mu\text{L}$) * | 14.60 ± 0.26 | 8.21 ± 0.35 | 4-12 | 16.26 | 0.000 |
| LYM (%) <sup>NS</sup> | 73.95± 1.727 | 45.73 ± 0 .821 | 40 -75 | 1.65 | 0.109 |
| MON (%) * | 7.51 ± 0. 20 | 3.55± 0.16 | 0-6 | 15.99 | 0.000 |
| NEU (%) * | 51.57± 1.54 | 33.54 ±0.91 | 10-50 | 10.38 | 0.000 |
| EOS (%) * | 15.06 ±0.35 | 6.72 ±0.39 | 0-10 | 15.53 | 0.000 |
| BAS (%) <sup>NC</sup> | 0.00 ± 0.00 | 0.00 ± 0.00 | 0-3 | - | - |

Reference range adopted from Sirois and Hendrix [55]. HGB = hemoglobin; HCT = hematocrit; MCV = mean corpuscular volume; MCH = mean corpuscular hemoglobin; MCHC = mean corpuscular hemoglobin concentration; RBC = red blood cell count; WBC = white blood cell count; LYM = lymphocytes; MON = monocytes; NEU = neutrophils; EOS = eosinophils; BAS = basophils. \*Significant at $p < 0.05$ ; NS = Non-Significant at $p < 0.05$ ; NC = No Change between groups.

### 3.2. Biochemical Findings

Regarding biochemical parameters, there were significant mean reductions (p < 0.05) in total protein, albumin, globulin, and the albumin-globulin ratio in the infected group (3.40 ± 0.10, 1.70 ± 0.07, 2.05 ± 0.07, and 0.86 ± 0.04, respectively) compared to the non-infected group (6.16 ± 0.04, 2.95 ± 0.04, 3.50 ± 0.11, and 0.86 ± 0.028, respectively). The biochemical parameters for non-infected sheep were within the established reference ranges (p < 0.05) (Table 2).

**Table 2.** Biochemical Parameters of *H. contortus*-Infected and Non-Infected Sheep.

| Parameters | Mean ± SE | | Normal Range | t-value | p-value |
| --- | --- | --- | --- | --- | --- |
|  | Infected (n=30) | Non-infected (n=30) |  |  |  |
| TP (g/dl) * | 3.40 ± 0.10 | 6.16 ± 0.04 | 6.0-7.9 | 21.22 | 0.000 |
| ALBU (g/dl) * | 1.70 ± 0.07 | 2.95 ± 0.04 | 2.4-3.9 | 20.04 | 0.000 |
| GLOB (g/dl) * | 2.05 ± 0.07 | 3.50 ± 0.11 | 3.5-5.7 | 10.20 | 0.000 |
| (AL/GL Ratio) * | 0.86 ± 0.04 | 0.86 ± 0.028 | 0.69-1.02 | 0.04 | 0.000 |

Reference range adopted from Sirois and Hendrix [55]. TP = Total Protein; ALBU = Albumin; GLOB = Globulin; AL/GL = Albumin/Globulin ratio. \*Significant at $p < 0.05$

### 3.2. Gross and Microscopic Lesions Characteriztion

#### 3.2.1. Gross lesions

When the abomasum was examined grossly, several important pathological findings were found. Focal ecchymotic hemorrhages, or localized areas of bleeding, were seen in the serosa (Figure 2A). There were *Haemonchus contortus* parasites and a nodular lesion in the paler, thickened fundic region (Figure 2B). Furthermore, there were *H. contortus* parasites, focal ulcerations, and paleness of the abomasal folds (Figure 2C).

**Figure 1.**
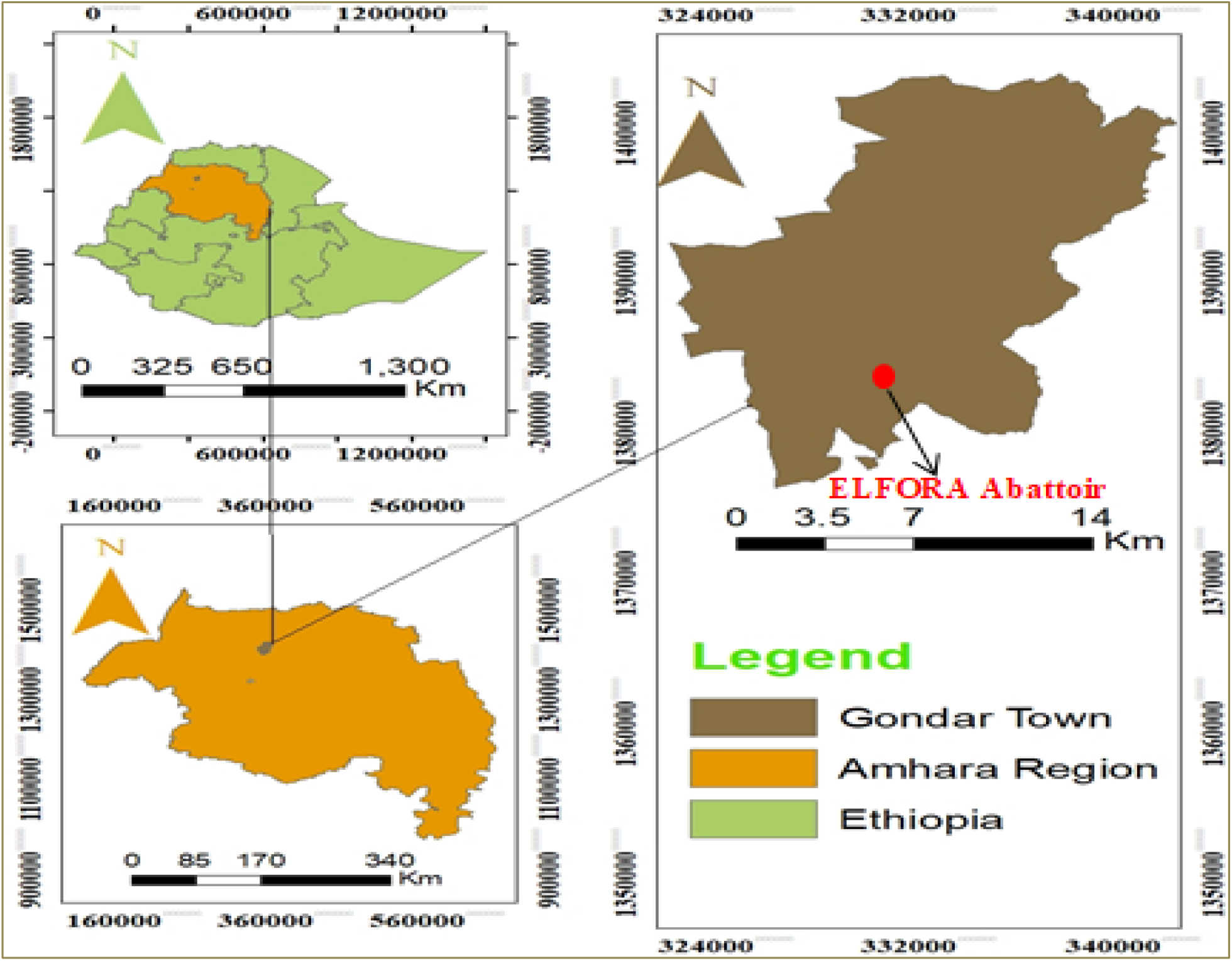
Map of the Study Area (ArcGIS 10.8)

**Figure 2.**
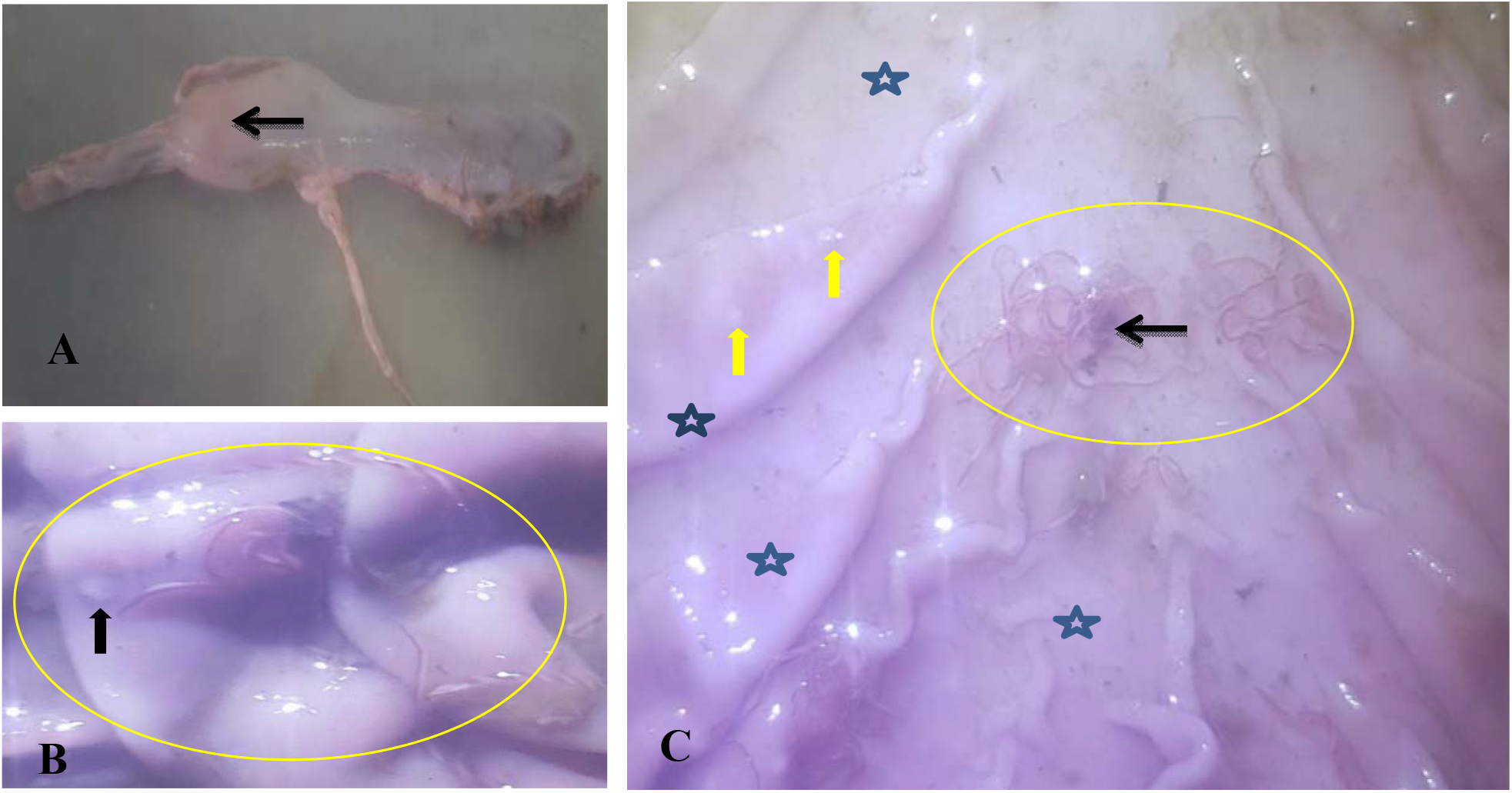
Gross Abomasum Lesions: A) focal ecchymotic hemorrhage in the abomasal serosa (arrow). B) A nodular lesion (black arrow) in the pale, thickened fundic region, and *Haemonchus contortus* parasites (circle). C) Pallor (star) was observed in the abomasal folds, along with *H. contortus* parasites (circle), nodular lesions (yellow arrows), and focal ulcerations (black arrows).

#### 3.2.2. Microscopic lesions

Microscopic examination revealed several notable pathological findings. The tissue sections demonstrated hemorrhage in the submucosa and the presence of tissue-dwelling worms along with inflammatory cell infiltration (Figure 3A). Furthermore, the mucosal glands were hyperplasia, and the collagen fibers contained clogged blood vessels (Figure 3B). There was a noticeable thickening of the muscularis mucosae and submucosa, as well as an abomasal gland hyperplastic proliferation (Figure 3C). In the fundic region, there was evidence of dilated abomasal glands, eosinophil infiltration, and denuded epithelium (Figure 3D). In addition, damage to the abomasum’s lamina propria and muscle layer was noted, along with submucosal hemorrhage. This was linked to constricted blood vessels and increased cellular infiltration in the submucosa and muscularis layers (Figure 3E). Lastly, inflammatory cell infiltration in the abomasal gland body and vacuolation in the fundic glands were demonstrated in a section displaying tissue-dwelling worms (Figure 3F).

**Figure 3.**
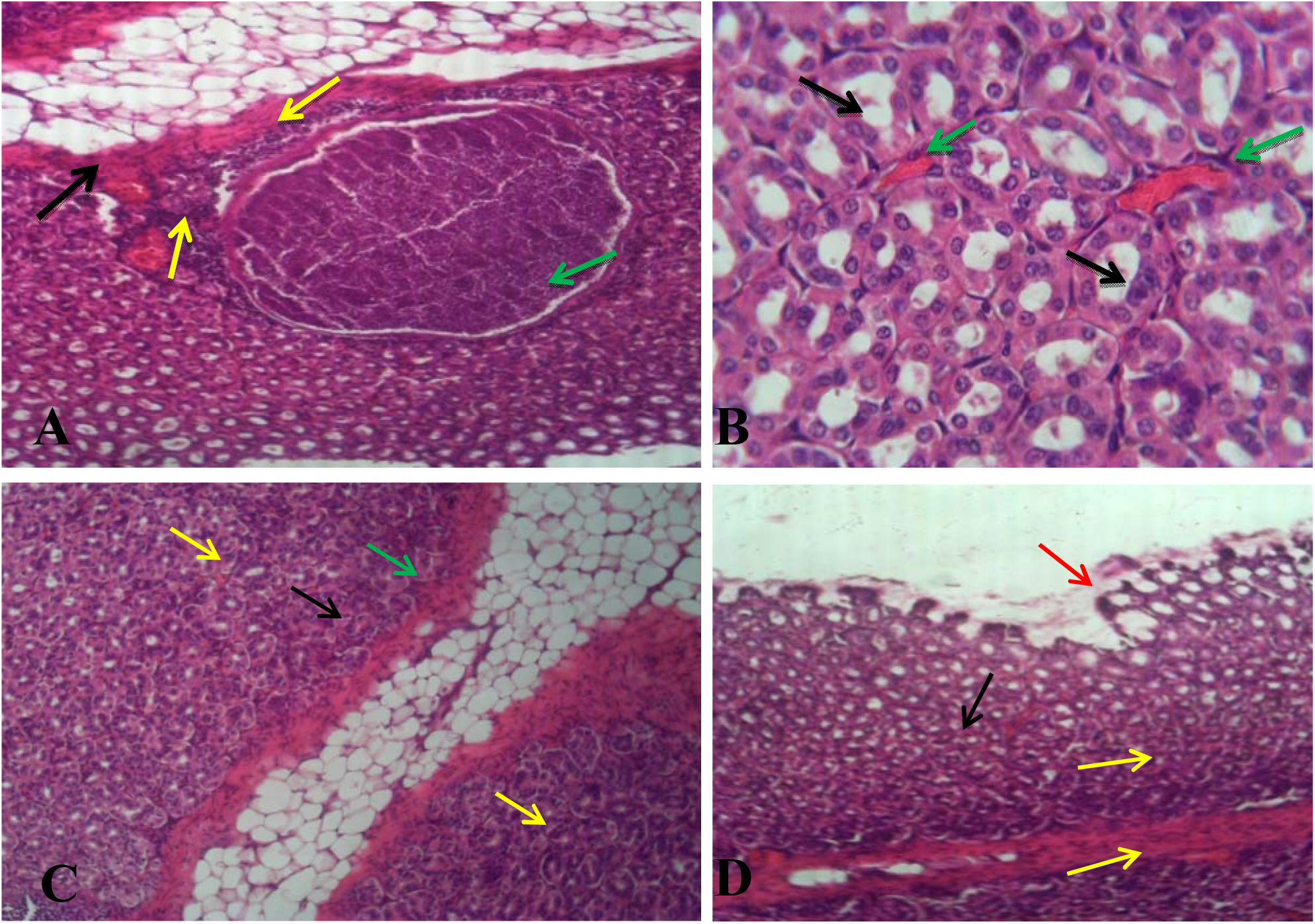
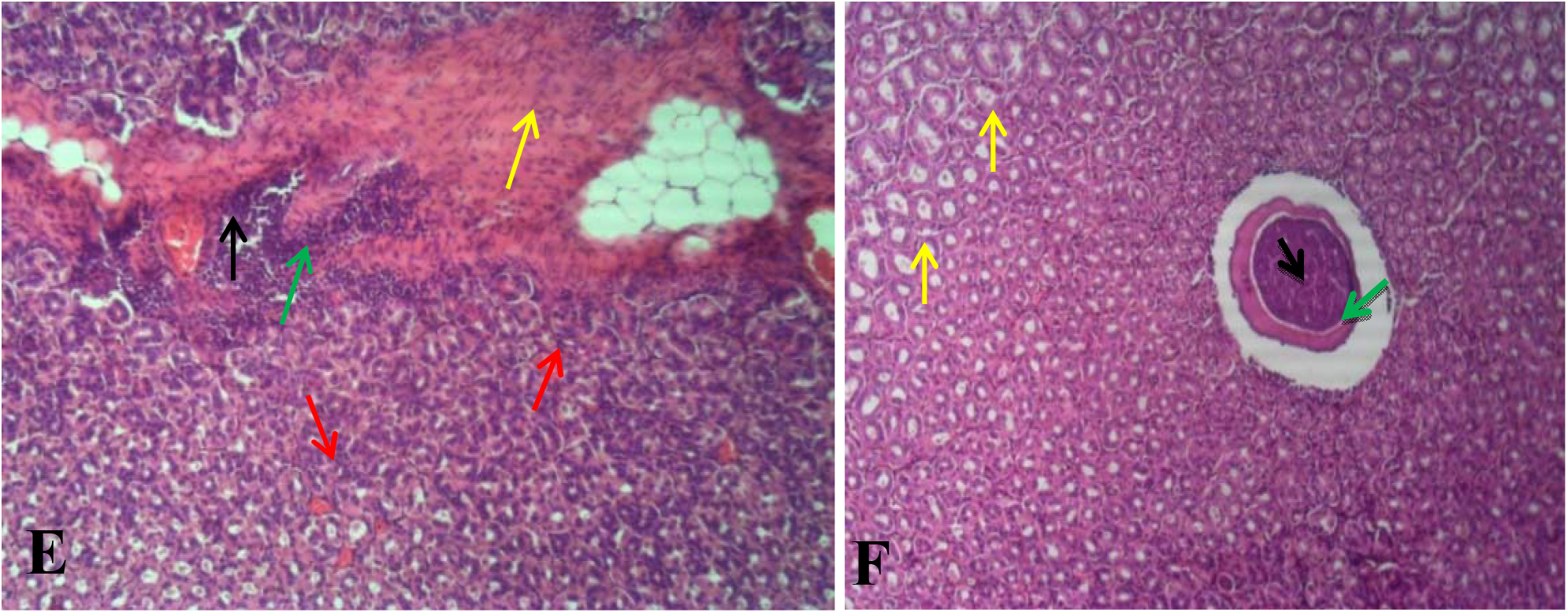
Microscopic Lesions of the Abomasum: A) Tissue-dwelling worm section (green arrow) with inflammatory cell infiltration (yellow arrows) and hemorrhagic submucosa (black arrow). H&E, 10X. B) Hyperplastic mucosal glands (black arrows) with congested blood vessels (green arrows) in the collagen fibers. H&E, 40X. C) Thickened muscularis mucosa (green arrow) and submucosa (black arrow), along with hyperplastic proliferation of abomasal glands (yellow arrows). H&E, 10X. D) Denuded epithelium (red arrow) with eosinophil infiltration (yellow arrows) and dilated abomasal glands (black arrow) in the fundic region. H&E, 10X. E) Hemorrhage in the submucosa (black arrow), damaged muscular layer and lamina propria of the abomasum (green arrow) with high cellular infiltration in the submucosa and muscularis layer (yellow arrows) and narrowed blood vessels (red arrows). H&E, 10X. F) Tissue-dwelling worm section (green arrow) with inflammatory cell infiltration in the body of the abomasal gland (black arrow) and vacuoles in fundic glands (yellow arrows). H&E, 10X.

## 4. DISCUTION

This study on ovine haemonchosis involved a comprehensive Haematobiochemical analysis and evaluation of pathological lesions. The infected sheep had significantly lower mean values of various parameters, including hematocrit (Hct), hemoglobin (Hgb), mean corpuscular volume (MCV), mean corpuscular hemoglobin (MCH), total erythrocyte count (TEC), total protein, albumin, globulin, and the albumin-to-globulin (AL/GL) ratio. Conversely, the total white blood cell counts (TWBC) and the counts of particular white blood cell types, such as neutrophils, monocytes, and eosinophils, were significantly higher (p < 0.05) in the infected sheep. There were no appreciable differences between sheep that were infected and those that were not in terms of the mean corpuscular hemoglobin concentration (MCHC) or lymphocyte counts; both were within the normal range. Furthermore, there was consistent similarity in the basophil counts between the two groups. The haemobiochemical parameters for non-infected sheep were within the established reference ranges. The parasite’s internal host disturbances may be the cause of the observed changes in hematological parameter values in infected sheep [21, 38]. On the other hand, variations in serum biochemistry might be linked to infection severity and tissue damage [27, 46]. The results of other researchers [12, 19, 24, 32, 35, 42, 47] were nearly in agreement with these findings.

Microcytic normochromic anemia is indicated by reductions in hemoglobin (Hgb), mean corpuscular volume (MCV), total erythrocyte count (TEC), hemoglobin (Hct), and mean corpuscular hemoglobin concentration (MCHC). A low MCV combined with normal MCHC values is the hallmark of this type of anemia, which may be the result of an iron deficiency brought on by significant blood loss from parasites. Although microcytic hypochromic anemia is the usual result of iron deficiency, it can also manifest at different stages as microcytic normochromic anemia [22, 59].

Other researchers [5, 9, 12, 19, 31, 36, 40, 42] also reported lower hemoglobin levels, hematocrit (Hct), and red blood cell (RBC) counts. These decreases were attributed to hemorrhage in the abomasum brought on by injuries from *Hemonchus contortus* infestations. On the other hand, increased values for the total white blood cell count (TWBC) and the counts of specific white blood cell types, such as neutrophils, monocytes, and eosinophils, were reported in the studies by Bakker et al. [14] and Irfan-ur-Rauf Tak et al. [34]. These increases are likely indicative of the immune response mounted by the host against the parasitic infection. Increased TWBC and certain types of white blood cells are indicative of a defensive response meant to neutralize the parasite and control the subsequent inflammatory response. Increased values were also reported for TWBC, eosinophils, and monocytes by Awad et al. [12].

A compensatory increase in eosinophils, neutrophils, and monocytes, all of which are crucial for the immune system’s reaction to parasitic infections, could account for the study’s normal lymphocyte percentage. This is in contrast to the results of Awad et al. [12], who found that infected sheep had higher lymphocyte counts. As observed in the current study, higher percentages of monocytes could be the result of monocytes serving as neutrophils’ backup line of defense [18]. On the other hand, infection-related stress may also be linked to an increased monocyte count [20]. Particularly in cases of parasitic infestation, elevated eosinophil counts are indicative of the body’s defense against the parasites [15, 53]. Given that both groups’ basophil counts were consistent, it is possible that allergic reactions, which are frequently mediated by basophils, do not play a major role in this parasite infection. Additionally, lower values for albumin, globulin, total protein, and the albumin-to-globulin (AL/GL) ratio were found in this study. These results suggest hypoproteinemia and hypoalbuminemia, which may be caused by a number of factors linked to parasitic infections. In particular, parasites can cause both an increase in protein loss and a decrease in protein synthesis. Further evidence for a relative increase in globulins is provided by the lower AL/GL ratio. Elevated inflammatory processes and the immune response are frequently linked to parasitic infestations [10]. Other researchers have reported similar findings of decreased total protein, albumin, globulin, and AL/GL ratio [12, 32, 35, 50].

Microscopic and gross lesions were common markers of abomasal pathology. On a gross level, this study revealed focal ecchymotic hemorrhages, or small patches of bleeding, in the serosa. In the paler, thickened fundic region, there was a nodular lesion and *Haemonchus contortus* parasites. Furthermore, there were *H. contortus* parasites, focal ulcerations, and pale abomasal folds. This result was also supported by earlier reports [1, 25, 35, 42, 48, 54].

The microscopic examination of the current study revealed infiltration of inflammatory cells, tissue-dwelling worms, and submucosal hemorrhage. In addition, there were blocked blood vessels in the collagen fibers and hyperplasia of the mucosal glands. The muscularis mucosae and submucosa showed discernible thickening, and there was also an abomasal gland hyperplastic proliferation. In the fundic region, there was evidence of dilated abomasal glands, eosinophil infiltration, and denuded epithelium. In addition to submucosal bleeding, there was damage to the abomasum’s muscle layer and lamina propria. This was linked to constricted blood vessels, vacuolation in the fundic glands, and increased cellular infiltration in the layers of the submucosa and muscularis. The results of this study were in line with earlier studies [1, 5, 7, 25, 35, 42, 48, 54].

A parasitic infection, particularly with *Haemonchus contortus*, may be indicated by tissue-dwelling worms, severe inflammatory cell infiltration, submucosal hemorrhage, hyperplasia of the mucosal glands, thickening of the muscularis mucosae and submucosa, hyperplastic proliferation of the abomasal glands, and clogged blood vessels embedded in the collagen fibers [8, 25, 28, 40, 44, 48, 49]. Denuded epithelium may have resulted from the potent chemicals released by activated inflammatory cells and the higher concentration of Haematophagus *H. contortus* in the abomasum [48]. The eosinophilic infiltration observed in the abomasal tissue may have been caused by an inflammatory response triggered by a *Haemonchus contortus* infection [16, 30, 52]. This infiltration may also be related to the host’s defenses against *H. contortus* [36, 39, 41, 57].

## 5. CONCULUTION

The current study was conducted to analyze the hematological and biochemical profiles of sheep infected with *H*.*contortus* along with those that are uninfected, as well as to identify the pathological lesions associated with this parasite. Infected sheep showed significantly elevated mean values of white blood cell counts and its types, particularly neutrophils, monocytes, and eosinophils. In contrast, most other haematobiochemical parameters showed significantly lower mean values. Grossly, ecchymotic hemorrhages, nodular formations, and *Haemonchus contortus* parasites were observed. Histopathologically, submucosal bleeding, mucosal gland hyperplasia, eosinophil infiltration, and tissue-dwelling worms were noted. The alterations in haematobiochemical parameters support the findings from both gross and microscopic lesions. Thus, integrating haematobiochemical analysis with the characterization of gross and microscopic lesions enhances the diagnosis of haemonchosis. Due to the significant hypoproteinemia observed, it is advisable to supplement helminth-infected animals with protein-rich feeds, such as legumes.

## Declarations

### Data sharing statement

Data is provided within the manuscript.

## Acknowledgements

The author expresses heartfelt gratitude to the staff of the College of Veterinary Medicine and Animal Science at the University of Gondar for their steadfast support. Special thanks are also extended to the personnel at the Gondar ELFORA slaughterhouse for their hospitality and invaluable assistance during the sample collection process.

## Author contributions

B.A.M: Conceptualization, Formal analysis, Visualization, Writing-original draft, Writing-review & editing; Investigation, Methodology, Funding acquisition, Validation, Software, Data curation, Resources. M.C.K & A.M.B: Supervision, Visualization, Validation. M.Y.M, M.Y, T.M & M.B: Visualization, Writing-review & editing, Validation. A.K & A.BT: Visualization, Writing-original draft, Writing-review & editing; Methodology, Validation, Resources.

## Ethics approval

Ethics approval for the study was granted by the University of Gondar’s College of Veterinary Medicine and Animal Sciences Ethics Research Review Committee (Ref. No. CVMASc/UoG/RERC/18/12/2023) on December 16, 2023. All animal procedures adhered to ethical guidelines, and verbal informed consent was obtained from the manager of Gondar ELFORA slaughterhouse, with the ethics committee’s prior approval before sample collection.

## Consent for publication

All authors have thoroughly reviewed and approved the final version of this manuscript. Each author has actively contributed to the work in various capacities.

